# Development of an efficient single-cell cloning and expansion strategy for genome edited induced pluripotent stem cells

**DOI:** 10.1101/2021.07.31.453934

**Authors:** Nupur Bhargava, Priya Thakur, Thulasi Priyadharshini Muruganandam, Shashank Jaitly, Pragya Gupta, Neelam Lohani, Sangam Giri Goswami, Saurabh Kumar Bhattacharya, Suman Jain, Sivaprakash Ramalingam

## Abstract

Disease-specific human induced pluripotent stem cells (hiPSCs) can be generated directly from individuals with known disease characteristics or alternatively be modified using genome editing approaches to introduce disease causing genetic mutations to study the biological response of those mutations. The genome editing procedure in hiPSCs is still inefficient, particularly when it comes to homology directed repair (HDR) of genetic mutations or targeted transgene insertion in the genome and single cell cloning of edited cells. In addition, genome editing processes also involve additional cellular stresses such as trouble with cell viability and genetic stability of hiPSCs. Therefore, efficient workflows are desired to increase genome editing application to hiPSC disease models and therapeutic applications. Apart from genome editing efficiency, hiPSC survival following single-cell cloning has proved to be challenging and has thus restricted the capability to easily isolate homogeneous clones from edited hiPSCs. To this end, we demonstrate an efficient workflow for feeder-free single cell clone generation and expansion in both CRISPR-mediated knock-out (KO) and knock-in (KI) hiPSC lines. Using StemFlex medium and CloneR supplement in conjunction with Matrigel cell culture matrix, we show that cell viability and expansion during single-cell cloning in edited and unedited cells is significantly enhanced. Our reliable single-cell cloning and expansion workflow did not affect the biology of the hiPSCs as the cells retained their growth and morphology, expression of various pluripotency markers and normal karyotype. This simplified and efficient workflow will allow for a new level of sophistication in generating hiPSC-based disease models to promote rapid advancement in basic research and also the development of novel cellular therapeutics.

## Introduction

Robust expansion capacity and high differentiation potential of human induced pluripotent stem cells (hiPSCs) have made for an invaluable tool for a vast array of biomedical and pharmaceutical applications including disease modelling, drug screening, toxicological assessment, gene therapy and development of cell based novel therapeutics (Müller and Lengerke, 2009; Maury et al., 2011; Rowe and Daley, 2019). Patient-derived hiPSC lines are intended to improve our understanding of patient heterogeneity, promote access to such model systems, and advance research into more effective regenerative therapies (Liu et al., 2019).

Recent advances in genome editing methods have significantly increased our ability to make precise alterations such as introducing disease causing genetic mutations or accurately correct patient derived hiPSCs (Govindan and Ramalingam, 2016). The combination of these two powerful technologies, enables the development of disease models to reach a new level of sophistication. Isogenic control and disease-specific hiPSC-lines have made it possible to model a variety of human diseases, which will overcome the issue of genomic variations between individual hiPSCs lines and also helps in gaining a better knowledge of their therapeutic potential (Wang et al 2014; Kim et al 2014; Kawatani et al 2021).

Conventionally, hiPSCs have been cultured in colonies on feeders and passaging as single cells is generally not suggested as this can result in spontaneous differentiation and genetic abnormalities (Dakhore et al., 2018). The challenge arises because hiPSCs are sensitive to various environmental cues such as osmolarity, pH, nutrition availability, mechanical stress, and most significantly, loss of cell-cell and cell-extracellular matrix (ECM) interaction (Mitalipova et al., 2005; Buzzard et al., 2004). Isolation of single cells and derivation of clonal cell populations with the necessary genetic alterations is one of the major bottlenecks of the genome editing procedure for pluripotent stem cells (Giuliano et al., 2019; DeWeirdt et al., 2020). Hence, the efficient selection and development of genome edited hiPSCs requires a single cell isolation approach paired with survival and improvement in expansion using optimal attachment aiding supplements and culture conditions.

It is well established that hiPSCs are difficult for single cell cloning. Traditional approaches such as manual picking of single colonies and limiting dilution cloning, are not ideal because they are time-consuming and inefficient (Singh, 2019; Vallone et al., 2019). Moreover, these approaches do not guarantee a successful single cell clonal expansion event after the isolation procedure. Genome editing procedures also subject stem cells to harsh environments such as electroporation, which significantly reduces their chances of survival (Chen and Miller, 2018). Hence, development of a simple and reliable procedure that enables robust and efficient single-cell derived clonal development of stable hiPSCs is highly desirable for research and clinical applications. In this report, we describe the detailed workflow for single cell cloning and expansion of clones of human control and genome-edited hiPSCs. Following the workflow detailed here, we demonstrate that single cell-derived hiPSC clones maintain the hallmarks of pluripotency and genomic stability.

## Materials and Methods

### Ethical approval

Ethical approval for the present study was obtained from the Ethics Committee (Ref no. CSIR/IGIB/IHEC/17-18/12) and Stem Cell Research Institutional Review Committee (Ref no. IGIB/IC-BT/8), CSIR-Institute of Genomics and Integrative Biology, New Delhi, India.

### Evaluation of different cell dissociation reagents

The hiPSC colonies from human control DYR0100 (ATCC; ACS-1011) and two patient-derived hiPSCs lines, IGIBi001-A (Bhargava et al., 2019) and IGIBi002-A (Thakur et al., 2021) were dissociated into single cells and plated at a density of 2500 cells per cm^2^ in 24 well dishes using different dissociation reagents, TrypLE, 0.5mM EDTA, ReleSR, Accutase and Gentle Cell Dissociation reagent (GCDR), in StemFlex medium supplemented with 1X RevitaCell. Media was changed every day and harvested for counting after 60 hours. To corroborate the viability in single cell attachment with different dissociation reagents, staining was performed with 500 µL of 0.25% crystal violet stain in 20% Methanol for 10 minutes at room temperature (RT). The cells were gently washed with Dulbecco’s Phosphate Saline Buffer (DPBS) 5-6 times to remove excess crystal violet stain and pictures were taken using a Cannon 550D DSLR camera.

### Synthetic guide RNA (sgRNA) and donor vector cloning

Chemically modified sgRNA targeting the T-cell receptor α constant *(TRAC)* locus (Eyquem et al., 2017) was used to edit hiPSCs. In order to achieve high frequency of editing and reduced off-target effects, 2’-O-methyl-3’-phosphorothioate modifications were included at the three terminal nucleotides of the 5’ and 3’ ends. Synthetic CAR and Turbo GFP under respective promoters were PCR amplified with 35 bp overhangs and assembled in pJHU3 vector using Gibson assembly mix as per the manufacturer’s instructions. The final vector pJHU3-CART-tGFP map is shown in Supplementary. Figure. 1.

### RNP preparation and Electroporation using Neon Transfection System

hiPSC lines of passage 10-20 were seeded at appropriate densities such that they were 65%-80% confluent on the day of electroporation. Prior to harvesting the hiPSCs for electroporation, the cells were pre-treated with 1X RevitaCell for an hour at 37°C. A fresh Matrigel coated 24-well plate with StemFlex medium and 1X RevitaCell was prepared and incubated at 37°C. For the KO experiment, one 10 µL Neon transfection reaction, 20 pmol of chemically modified *TRAC* sgRNA and 1µg Cas9 was incubated at RT for 15 minutes. Electroporation was carried out with 2 × 10^5^ cells in Neon Buffer R with RNP complex at 1200V, 30 msec, 1 pulse. For the KI experiment, 2.0 µg of CART donor plasmid was also added to the electroporation reaction containing the aforementioned Cas9-sgRNA RNP complex. The electroporated cells were added to previously prepared Matrigel coated 12-well plate containing fresh culturing medium supplemented with 1X RevitaCell. The medium was changed within 4-6 hour post-electroporation without RevitaCell supplement and cells were allowed to recover for another 48-72 hours.

### FACS Sorting

Bulk Sorting: In case of KI experiment, the electroporated hiPS cells were sorted based on GFP marker present in the donor construct, 48 hours post-electroporation, and expanded before proceeding to further experiments. The GFP expression was transitory and hence, sorting enriched the electroporated cell population.

Single Cell Sorting: hiPSCs were sorted when they were 60-80% confluent, or 72 hours post electroporation in case of KO experiment. On the day of sorting (Day 0), cells were pre-conditioned with 1X RevitaCell, 1 hour before harvesting for sorting. After 1 hour they were harvested using Accutase and a single cell suspension was made. The cells were resuspended in 3-5% Knockout Serum Replacement (KOSR) in DPBS after one spin wash with DPBS. The cells were sorted in 96 well plates coated with Matrigel containing 100ul of cloning Media (StemFlex supplemented with 1X CloneR or 1X RevitaCell). The experimental conditions for single cell clone expansion with Revita Cell and CloneR are described below.

CloneR Supplement: The media was replaced completely with fresh cloning media on Day 2. Additional cloning media of 50 µL was added to the wells on Day 3. From then onwards 50 µL media was replaced with fresh culturing media (Stem Flex without 1X CloneR) every alternate day till Day 7. Thereafter full media changes were performed for wells with positive clones which emerge around day 3-5 and are ready to be harvested around day 10-12.

RevitaCell Supplement: The media was replaced completely with fresh cloning media on Day 3. Additional cloning media of 50 µL was added to the wells on Day 6. From then onwards 50 µL media was replaced with fresh cloning media (StemFlex supplemented with 1X RevitaCell) every two days till Day 9. Thereafter full media changes were performed for wells with positive clones which emerge around day 5-7 and are ready to be harvested around day 12-15.

### Screening of gene edited single cell clones

Clones from 96 well plates were transferred to 24 well plates and were taken for genotyping analysis. Screening was done by making cell lysate of clones for which harvested cells were centrifuged at 1000 rpm for 5 mins. Following one DPBS wash, cells were resuspended in 30µL 5mM Tris-HCl and incubated at 95°C for 15 min. The reaction was allowed to cool at RT and then Proteinase K was added and further incubated at 56° for 30 min and 85° for 5 min. This cell lysate was allowed to cool at RT and was used for screening for KO and KI clones. To analyse the KO clones, we performed T7 Endonuclease 1 assay following the manufacturer’s instructions. To analyse the KI clones, we performed 5 ‘junction and 3’ junction PCR analysis using cell lysate prepared from single cell clones. List of primers and PCR conditions used are mentioned in Supplementary Table No. 1

### Immunostaining of single cell derived hiPSC clones

The immunostaining was performed by fixing the cells using 4% paraformaldehyde at RT for 15 min, followed by 1% Triton X-100 permeabilization for 10 min at RT. A blocking solution (5% bovine serum albumin) was added to the cells and the cells and were incubated for 30 min at RT. Staining was carried out with appropriate primary antibodies diluted in PBS, cells were incubated for 12-16 hours at 4°C followed by respective secondary antibodies for 1 hour at RT. A 15-min staining procedure with DAPI was performed for nuclear staining. Using a fluorescence microscopy station (Floid cell imaging station), cells were visualized, and images were taken. The antibody details are listed in Supplementary Table No.2.

### Karyotyping analysis

G-banded karyotyping was used to analyze the genetic stability of randomly selected single cell derived hiPSC clone. Actively dividing hiPSCs at the metaphase stage were arrested with colcemid for 25 min at 37°C, followed by dissociation. Hypotonic potassium chloride was applied to the cells, and they were then fixed in acetic acid and methanol (3:1 v/v) solution overnight, and further air dried. GTG banding was performed on metaphase spread slides. Leica DM2500 was used to acquire images, and Cytovision V7.7 program was used for analysis.

### Statistical analysis

We have performed all experiments in replicates of two (n=2). The statistical analysis and graphs were made using Prism9 Graph Pad. A p value of < 0.05 was taken into account for determining statistical significance. Schematics were prepared in Biorender (“BioRender”, 2021). Vector map was generated using SnapGene.

## Results

### 1. Cell dissociation solution Accutase supports maximum cell recovery of single cell passaged hiPSCs

Single cell passaging is a foundation for many downstream applications such as cell sorting, genome editing, imaging, single cell cloning, high-throughput drug screening and directed differentiation of certain lineages. It is widely accepted that single-cell passaging is generally accompanied by significant loss of cell viability. For routine hiPSC dissociation, both enzymatic and enzyme-free dissociation reagents have been widely employed. Elements that might increase single-cell survival after FACS sorting have then been investigated to see how the most effective single cell cloning can be accomplished under a feeder-free environment (Figure 1). In order to evaluate which dissociation reagent is more suitable for single cell passaging and overall cell survival, we tested different commercially available dissociation reagents TrPLE, 0.5mM EDTA, ReLESR, Accutase and Gentle Cell Dissociation Reagent (GCDR) (Figure 2A) Post passaging, cells were allowed to recover for 24 hours in StemFlex medium supplemented with 1X RevitaCell, followed by daily media change with StemFlex culturing medium alone. Cell survival and recovery was monitored using phase-contrast microscopy. We observed maximum attachment and cell survival with very minimum cell death with Accutase followed by GCDR. The same trend was observed when cells were harvested and counted 60 hours post-passaging (Figure 2B). This result was consistent with crystal violet staining indicative of live and attached hiPSCs (Figure 2C). Taken together, our findings suggest that Accutase supports routine single cell passaging of hiPSCs and was thus used for our subsequent experiments.

**Figure 1.**
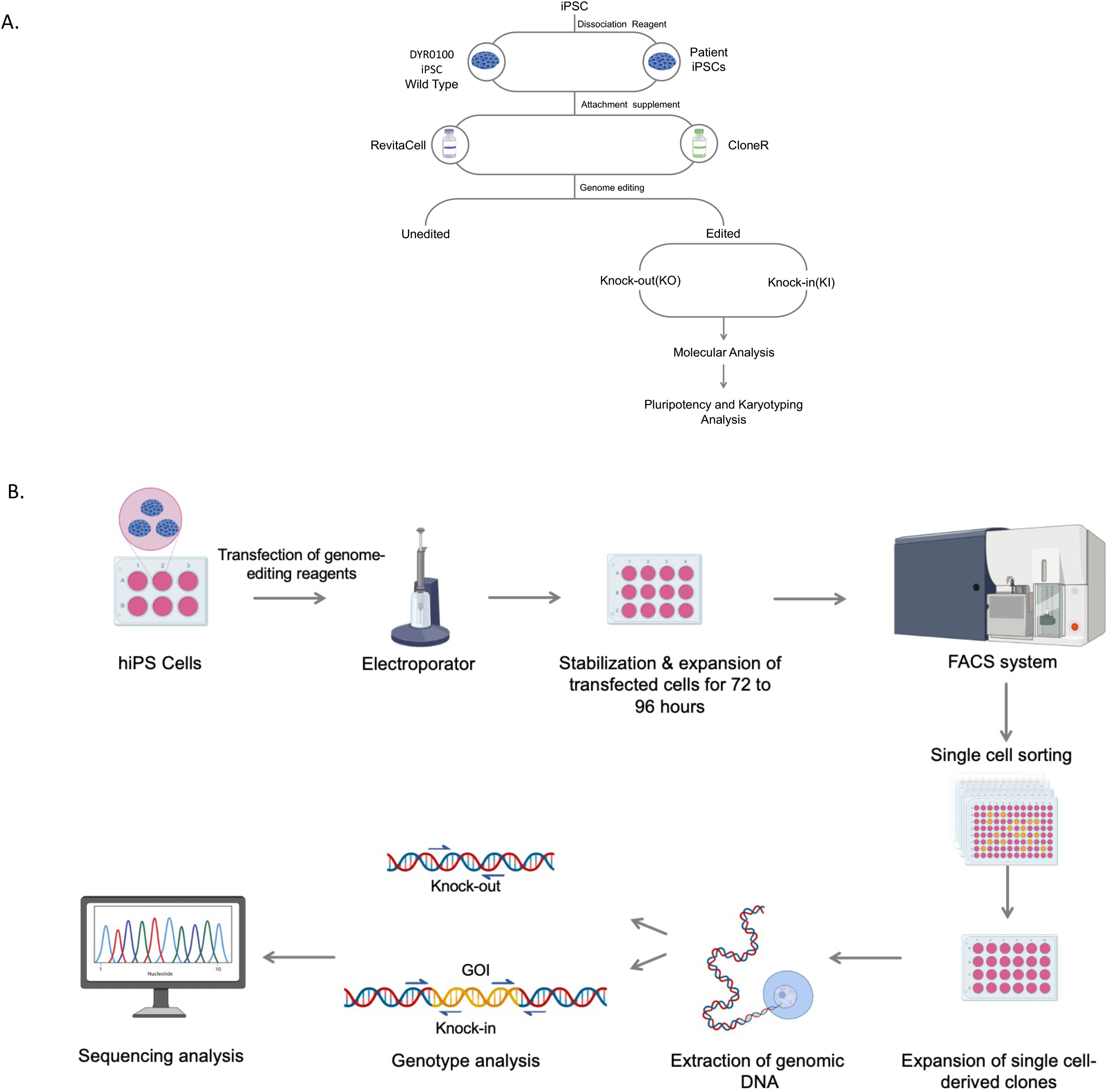
Schematics showing development of an efficient single cell cloning strategy for genome edited hiPSCs.(A) Overview of the single-cell cloning strategy. The strategy was tested with different hiPSC lines before and after genome editing. Two different attachment aiding reagents, CloneR and RevitaCell were compared for single cell cloning efficiency with edited and unedited hiPSC lines. Randomly selected single cell derived clones were tested for their pluripotency and chromosomal integrity. (B) Workflow showing the nucleofection of hiPSC lines with genome-editing reagents followed by FACS sorting in 96-well plates as one cell/well with above-mentioned aiding reagents. Subsequently, the single cell-derived genome-edited clones were subjected to molecular analysis.

**Figure 2.**
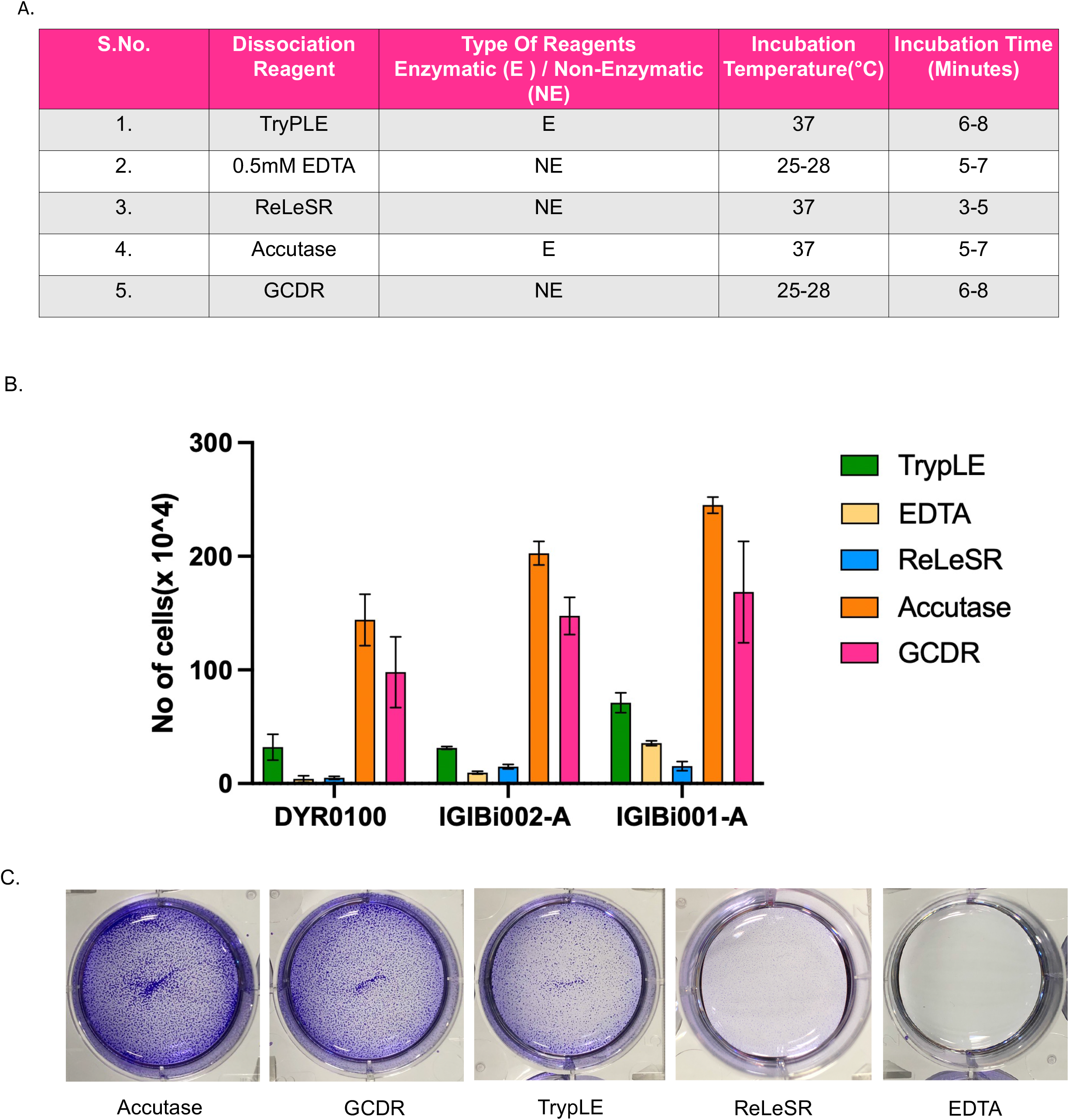
Assessment of different cell dissociation reagents for single cell attachment .(A) Different enzymatic (E) and non-enzymatic (NE) passaging reagents and their dissociation conditions. (B). Comparison of different dissociation solutions for single cell passaging. The cells were dissociated and seeded at same density and harvested and counted 60 hours post-passaging(n=2). (C) Crystal violet staining indicative of live and attached hiPSCs.Error bars denote the mean ± SD, p< 0.05.

### 2. Successful Single cell clone generation from edited and unedited hiPSC lines

We compared two attachment aiding supplements, CloneR and RevitaCell for single cell cloning of hiPSCs (Figure 3A). To achieve reliable high-throughput isolation of single cell-derived hiPSC clones in 96-well plates using FACS, we first established a stringent gating approach to ensure sorting of healthy and single cells (Figure 3B). During the initial days when the single cell is expanding to form a colony, the morphology may not be ES like but more elongated (Figure 3C). We observed colony formation as soon as 3-5 days after single cell sorting. These cells compact over time and eventually depict the typical morphological characteristics of hiPSCs (Figure 3C). We obtained approximately 10-15% more single cell clones when hiPSCs were cultured with CloneR than with RevitaCell across edited and non-edited hiPSC single cell cloning experiments (Figure 3D). The quality and health of hiPSC clones culture with both supplements were consistent and comparable. We took a threshold for determining clonal efficiency at Day 12 post sorting. We observed that the cells cultured with RevitaCell are relatively slower growing than with CloneR. By extending the exposure window of CloneR or RevitaCell, we were able to enhance the clone survival significantly.

**Figure 3.**
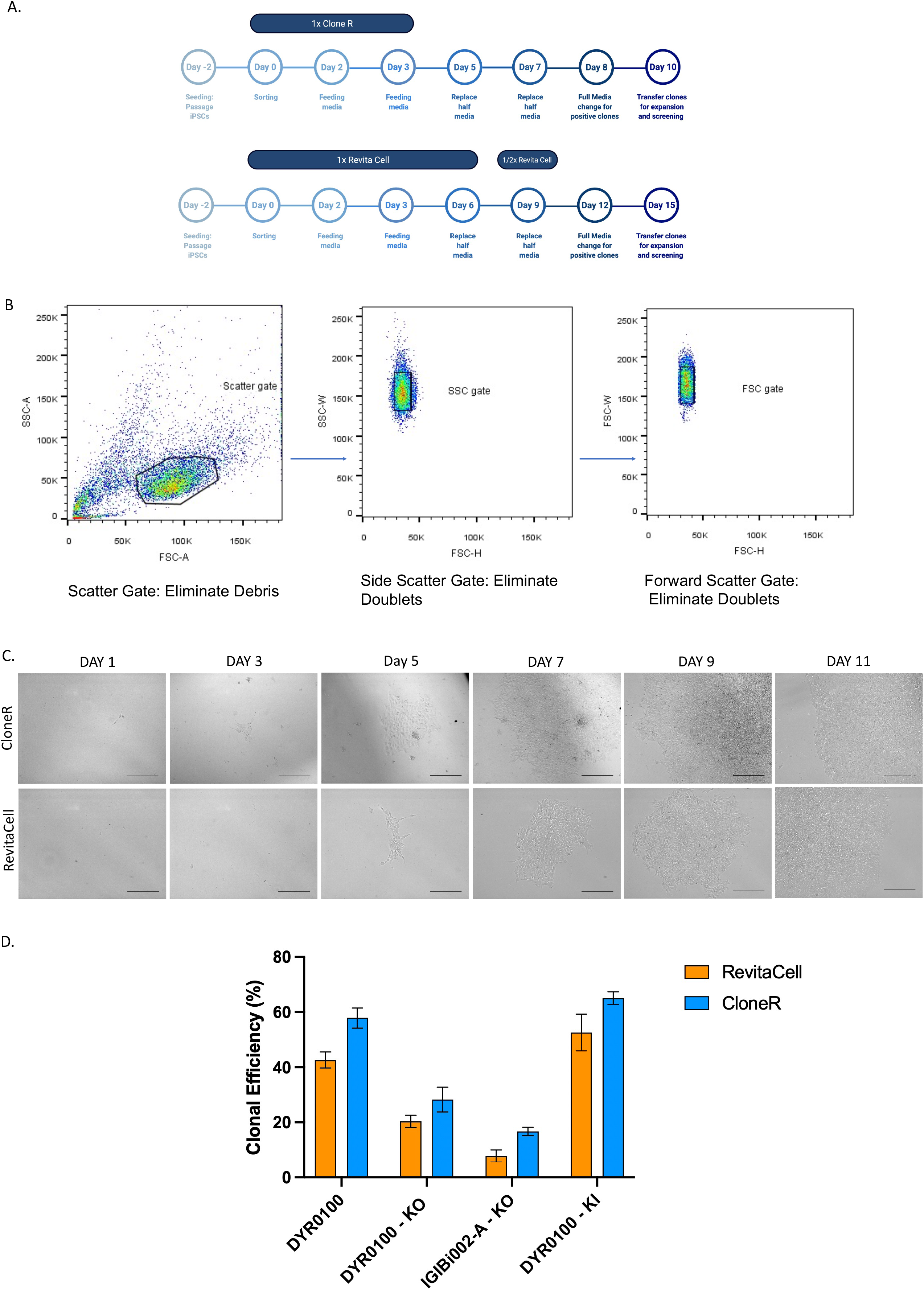
Efficient single cell cloning in hiPSCs. (A) Methodology overview for achieving single cell hiPSC clonals. (B) Representative FACS gating for hiPSC sorting.(C) Phase contrast images were taken at 48 hours intervals depicting emergence of single hiPSC clones till Day 12. Images depict the development of a healthy colony arising from a FACS sorted single cell. (D) Analysis of clonal efficiency comparing different attachment supplements(n=2). Error bars denote the mean ± SD, p< 0.05.

### 3. Single-cell cloning workflow successfully produced KO and KI clones

The genome editing strategy we employed to demonstrate the efficiency of our single cell cloning workflow for hiPSCs is shown in Figure 4A and Figure 4D. We performed electroporation to deliver Cas9-sgRNA RNP complex in hiPSCs and thereafter went ahead with scatter plot-based sorting of the cells after 72 hours of electroporation. We determined the cleavage efficiency of the sgRNA with T7 endonuclease 1 assay in the heterogeneous cells to confirm that the Double strand break (DSB) had occurred (Figure 4B). Positive single clones were expanded for further screening of edited clones. We performed T7 endonuclease 1 assay to screen the single-cell KO clones and achieved high efficiency of approximately 50% KO events in all hiPSC lines tested (Figure 4C). For the purposes of our study, we have considered T7 endonuclease I positive clones as “knock-out clones”. These clones could have heterozygous or homozygous indels caused due to NHEJ. Depending on the nature of the gene editing strategy, further confirmation of the KO clones can be made through sequencing analysis.

**Figure 4.**
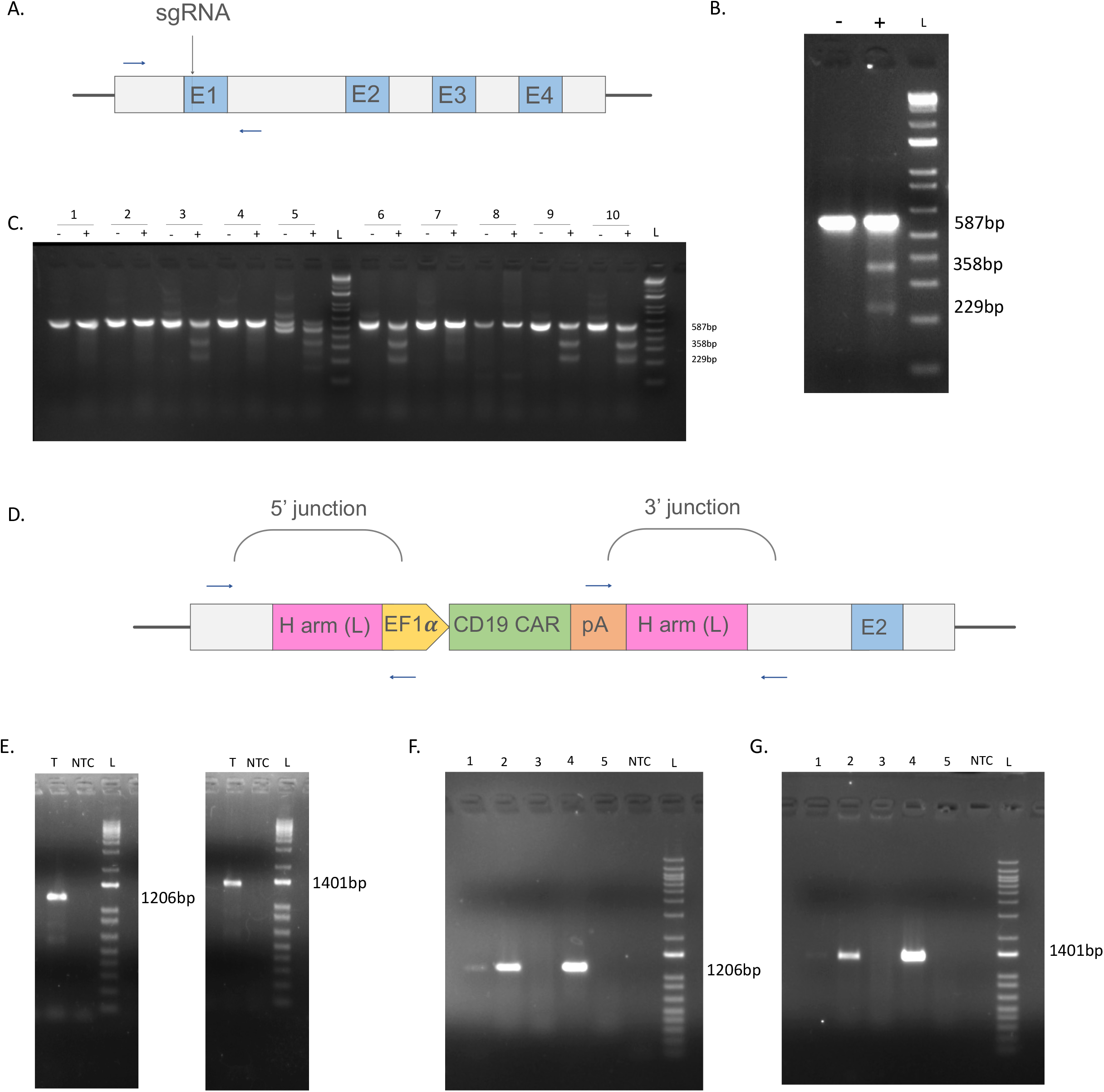
Screening of gene edited single cell derived hiPSC clones.(A) Schematic showing the sgRNA target site for double stranded break in the *TRAC* locus and the primers designed to detect cleavage efficiency. T7 Endonuclease I assay for (B) heterogeneous electroporated hiPSCs for KO strategy (- : PCR product without T7 Endonuclease 1, + : PCR product with T7 Endonuclease 1) and (C) single cell derived hiPSC clones (- : PCR product without T7 Endonuclease 1, + : PCR with T7 Endonuclease 1 enzyme). (D) Schematic depicting screening primers of the 5’ and 3’ junctions of the donor DNA insertion sites.(E) Screening and confirmation of KI with junction PCR at 5’ and 3’ KI site in GFP sorted population (T: template control, NTC: no template control) . (F) 5’ junction analysis of donor integration of single cell derived hiPSC clones (T: template control, NTC: no template control). (G) 3’ junction analysis of donor integration of single cell derived hiPSC clones. The expected size for amplicons for 5’ junction is 1.2 kb and for 3’ junction is 1.4 kb for site-specific insertion of the donor at the targeted *TRAC* locus in the genome. (T: template control, NTC: no template control) (L) is 1kb+ ladder.

For our KI strategy we performed electroporation with Cas9-sgRNA complex along with a donor vector. We sorted the cells based on a GFP marker in the donor vector 48 hours post electroporation. The GFP positive sorted cells were thereafter expanded before proceeding to single cell sorting. We confirmed that the KI has occurred by performing Junction PCR at the insertion site in the GFP enriched sorted population (Figure 4E). In order to analyze the positive single-cell derived KI clones, we performed PCR analysis of the 5’ and 3’ junctions of the donor DNA insertion sites using one of the primers anchored outside the *TRAC* homology arms of the donor and the other anchored within the donor DNA. KI single-cell clones yielded the expected 1200bp (5’ junction) and 1400bp (3’ junction) fragments, confirming donor insertion at the *TRAC* locus in the hiPSCs (Figure 4F and Figure 4G). Around 15% of the screened clones amplified for both junctions. We furthermore sequenced the junction PCR amplicons to determine the nucleotide sequence at the 5’ and 3’ junctions of the donor insertion site in the CART hiPSCs. Sequencing results further confirmed the presence of CAR donor DNA at *TRAC* locus resulting from homologous recombination (Supplementary Figure 2).

### 4. Single-cell derived clones retained morphology, expansion, pluripotency and normal karyotype

It is important to make sure that the genome-edited single-cell derived hiPSC clones show typical embryonic stem cell (ES)-like colonies with compact cells and clean borders, as well as a high nucleocytoplasmic ratio. Moreover, these cells should be expandable with no differentiation. In order to test this, we have passaged edited single-cell derived clones for at least 4 passages and also freeze-thawed the edited clones. In both the cases, clones demonstrated pluripotent identity with no differentiation (Figure 5A). Expression of pluripotency markers is a strong indicator of their stemness. Hence, we have shown the expression of pluripotent markers OCT4, SOX2, SSEA4 and TRA-1-60 by immunostaining after single-cell cloning and expansion (Figure 5B). Furthermore, we evaluated chromosomal stability of edited single-cell derived clones. A representative chromosomal karyotyping analysis of an expanded clone revealed a normal karyotype (Figure 5C), without any visual aberrations. Overall, our single-cell cloning workflow can support the viability, expansion and maintenance of pluripotency following genome editing procedure.

**Figure 5.**
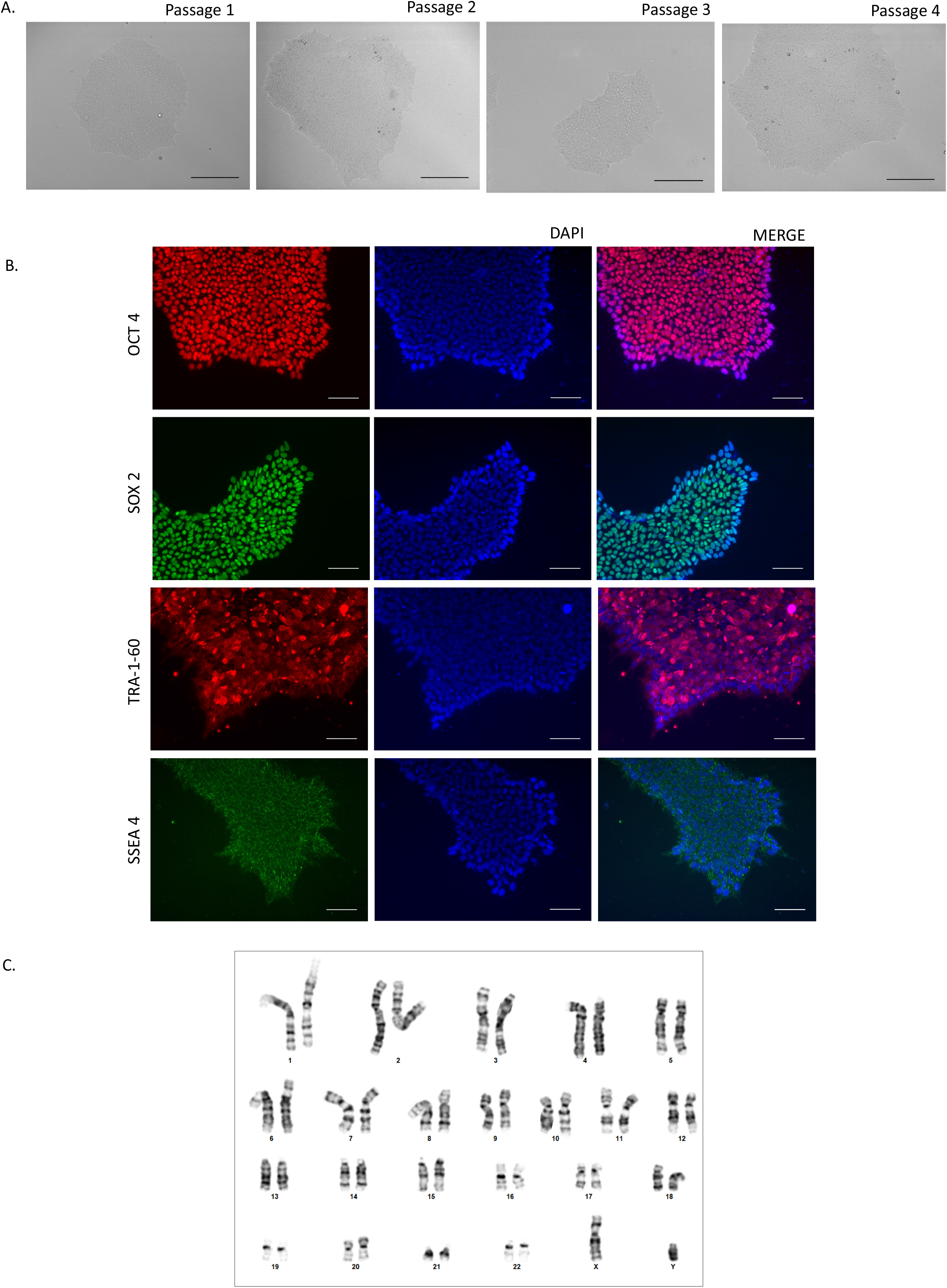
Characterization of a single cell derived edited hiPSC clone.(A) Phase contrast images showing maintenance of hiPSC morphology in an edited clone through multiple passaging.(B) Immunofluorescence staining for pluripotency markers,OCT4, SOX2, SSEA4 and TRA-1-60 after single-cell cloning and expansion in an edited hiPSC clone. For nuclei staining, DAPI was counterstained. Scale bar, 100µm (C) Genetic stability of the single-cell sorted hiPSC clones was investigated by G-banded karyotyping analysis. A representative karyotype of the edited hiPSC clone showing normal XY karyogrm post editing and sorting.

## Discussion

Recent advancements in genome editing have allowed the introduction of targeted genetic modifications in human cells and organisms (Salsman and Dellaire, 2017; Wang et al., 2016; Adli, 2018; Manghwar et al.,2019). These methods have also been employed in hiPSCs for functional studies of genetic diversity as disease models, and in the broader field of regenerative medicine. However, a considerable obstacle still exists in achieving single-cell cloning in gene edited hiPSCs (Reubinoff et al., 2000; Pyle et al., 2006; Amit et al., 2000). It is currently challenging to handle isolated clonal cells in a way that allows them to survive and thrive during genome editing (Frisch and Screaton, 2001). Because delicate hiPSCs grow in colony form and need a feeder-dependent environment, re-plating as single-cells eliminates growth cues and survival, thus limiting viability. For the expansion of any clonal colonies carrying the genetic mutation of interest from the genome editing procedure, promoting survival and proliferation at the single-cell level is crucial. The isolation of single cell clonal lines, which is normally a labor-intensive process, is one of the key bottlenecks of achieving an efficient genome-editing strategy that we intended to overcome with this study.

Though a couple of studies have shown methods for clonal expansion of genome edited pluripotent stem cells, these methods rely largely on feeder-dependent culture system, manual colony picking, limited dilution, and drug selection (Singh, 2019). The above approaches are both laborious and time-consuming and there is no actual proof of clonality. Hence, there is still a lack of standard workflow for hiPSC single cell cloning of genome edited cells.

Recently Singh, 2019 has reported single cell cloning protocol for human pluripotent stem cells using a feeder dependent environment which employs irradiated mouse embryonic fibroblasts (iMEF) (Singh, 2019). Though iMEFs supports long-term growth of hiPSCs, the feeder cell layer (iMEF) presents a number of concerns, feeder cells are susceptible to batch-to-batch variability, and it may secrete unknown components into the culture media, which may cause xenogenic contamination in the culture system (Higuchi et al., 2015). In addition, earlier reports indicate that cultivation of hESCs may have negative impacts as retroviral contamination may exist in iMEF (Cobo et al., 2008). Thus, the clinical application of hiPSCs can be hindered by xenogenic components. In the last several years, numerous evidence indicates that both the physical signals and biological signals from the culture materials including basement membrane, direct stem cell fate during growth and maintenance of their pluripotency (Chermnykh et al., 2018; Costa et al., 2012).

Single cell cloning with hiPSCs is still a big hurdle even in a feeder independent environment as they have low survival and attachment when dissociated as single cells (Chen et al., 2014). Hence, we aimed to develop an efficient workflow for single cell cloning of hiPSCs. In regular maintenance of hiPSCs, they are dissociated as aggregates with a gentle and non-enzymatic reagent (Vazin and Freed, 2010). First, we set out to compare which dissociating reagent results in maximum attachment of viable hiPSCs when passaged as single cells. Towards this objective, we found accutase performed best among other cell dissociation reagents in terms of cell survival, expansion and resulted in minimal cell death when cultured in StemFlex medium supplemented with 1X RevitaCell. This result is consistent with previous reports which have shown that Accutase does not affect the single cell viability and growth rate of pluripotent stem cells (Bajpai et al., 2008). Furthermore, an attachment supplement helps in reducing the impact of stress from single cell passaging and significantly enhances overall cell viability and survival (Dakhore et al., 2018).

Subsequently, we proceeded with Accutase for dissociation of hiPSCs and went on to compare which attachment aiding supplement works efficiently for single cell cloning. We investigated this in different hiPSC lines, both in non-edited as well as edited cells. We found CloneR to consistently perform better than RevitaCell, resulting in 10%-15% more clones. We have shown that in a KO gene editing strategy, the trend of our results is maintained in both wild type as well patient derived hiPSC lines. However, we overall recovered less number of clones compared to non-edited DYR0100, in KO strategy as cells were sorted 72 hours post electroporation, without allowing them to recover completely from stress of electroporation. We reasoned that for KO strategy, it may not be essential to enrich the population as NHEJ is the default repair mechanism that occurs throughout the cell cycle (Currall et al., 2013) and even in a heterogeneous pool, there is a high chance of getting a KO clone due to creation of indels. Since the cleavage efficiency of our sgRNA was good, as confirmed with a T7 endonuclease 1 assay in a heterogenous population of electroporated cells, we had proceeded to single cell sorting without further enriching the population. During screening of scatter plot based sorted clones, we achieved almost 50% positive KO clones. Hence this workflow was suitable for our KO strategy, if however, the cleavage efficiency of sgRNA is low then enriching the population prior to single cell sorting is recommended. Our findings suggest that single cell sorted clones, cultured with CloneR emerge earlier than when cultured with RevitaCell and are ready to be passaged at Day 10-12 post single cell sorting.

We proceeded with carrying out a KI experiment in our target cell line, DYR0100. As it is known, homology directed repair (HDR) only occurs in the G1-S phase of the cell cycle thus achieving single cell cloning with a KI strategy is less efficient (Yang et al., 2020). Hence, we enriched the KI edited population by sorting the hiPSCs post 48 hours of electroporation, based on a GFP marker in the donor construct which expresses transiently. The GFP positive sorted hiPSCs were then expanded and stabilized before proceeding to single cell sorting. The number of clones we recovered with our single cell cloning workflow was comparable to the scatter plot based sorted non edited stable hiPSC line DYR0100. Similar trend was observed in respect to attachment factors. We have also observed that directly going for single cell sorting based on marker expression without enriching the population beforehand yields less clones (data not shown). Hence, we recommend, enriching the cell population that has taken up the donor construct based on a fluorescent marker and then going for scatter plot-based sorting.

Together, we found CloneR to perform consistently better than RevitaCell in aiding attachment and expansion of single cell derived clones in both edited and non-edited hiPSCs. The attachment factor did not affect the number of positive edited clones recovered. The higher clonal efficiency in KO clones was observed in comparison to KI experiments which is consistent with previous reports when such strategies are adopted (Niccheri et al., 2017; Wang et al., 2012; Xie et al., 2017).

Additionally, we have also tested the stemness and proliferation of randomly selected single cell derived clones after freeze-thawing. We observed cells recovered within 24 hours post thawing and no change in morphology or growth was observed. Characterization of single cell derived clones expressed pluripotent markers and did not show any chromosomal abnormalities. Most significantly, we show that this strategy enables efficient single cell cloning for genome edited hiPSCs. Combining these technologies and our single cell cloning workflow led to greater success in the generation of homogenous genome-edited hiPSC clones and offer novel approaches to study disease biology *in vitro*.

## Summary

We developed a reliable and efficient approach that supports single-cell cloning and expansion of edited (both KO and KI) hiPSCs, with generated clones retaining normal ES-like morphology, proliferation and pluripotent markers expression, and making it an ideal system for accelerating experiments that need sensitive handling of genome-edited hiPSCs. This efficient method should contribute to a diversity of applications in human disease biology, including generation of isogenic hiPSC-derived disease modeling for basic research, development of cellular therapeutics and high-throughput drug screening.

## Acknowledgements

This work was supported by the Early Career Award, Science and Engineering Research Board (SERB) (Grant No. ECR/2017/002212) to SR, Department of Biotechnology grants (BT/GET/119/SP31652/2020 and BT/GET/119/SP31652/2020) Government of India (GoI) to SR. NB is a recipient of Senior Research Fellowship from Indian Council of Medical Research, Government of India. PT, PG and SGG are recipients of Senior Research Fellowship from Department of Biotechnology, Council of Scientific and Industrial Research and University Grant Commission, GoI respectively. We would like to thank Dr. Deepak Rathore of Translational Health Science and Technology Institute (THSTI) for his technical assistance in FACS sorting experiments.

## Author’s contribution statement

S.R conceived and supervised the experiments. S.R, N.B, P.T designed the experiments. S.J and T.P helped with developing hiPSC lines. N.B, P.T, T.P, S.J, P.G, S.B, N.L and S.G.G carried out experiments. N.B, P.T and S.R analyzed the data and wrote the manuscript.

---

**Table 1.**
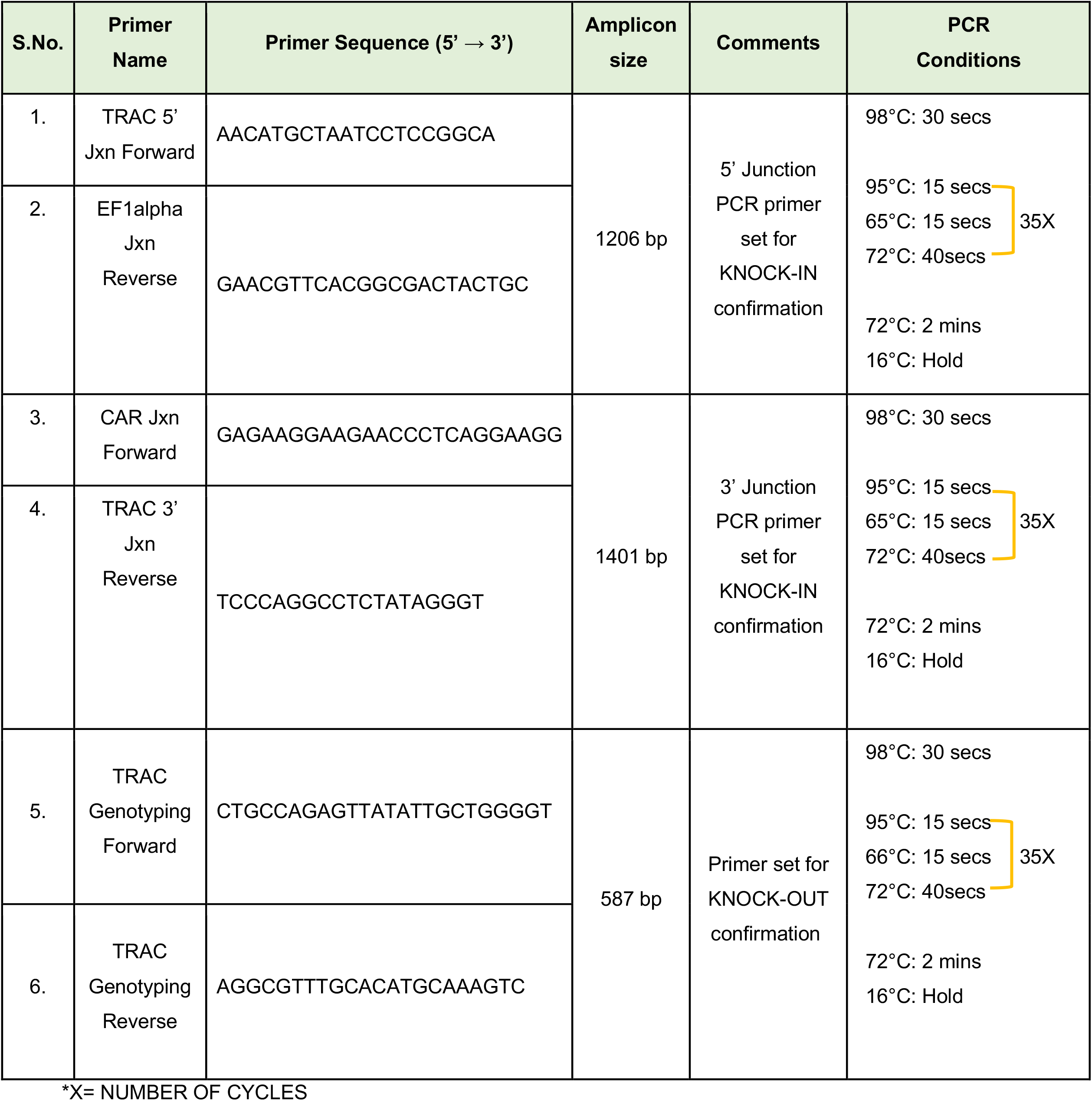
Primer Details and PCR Conditions.

**Table 2.**
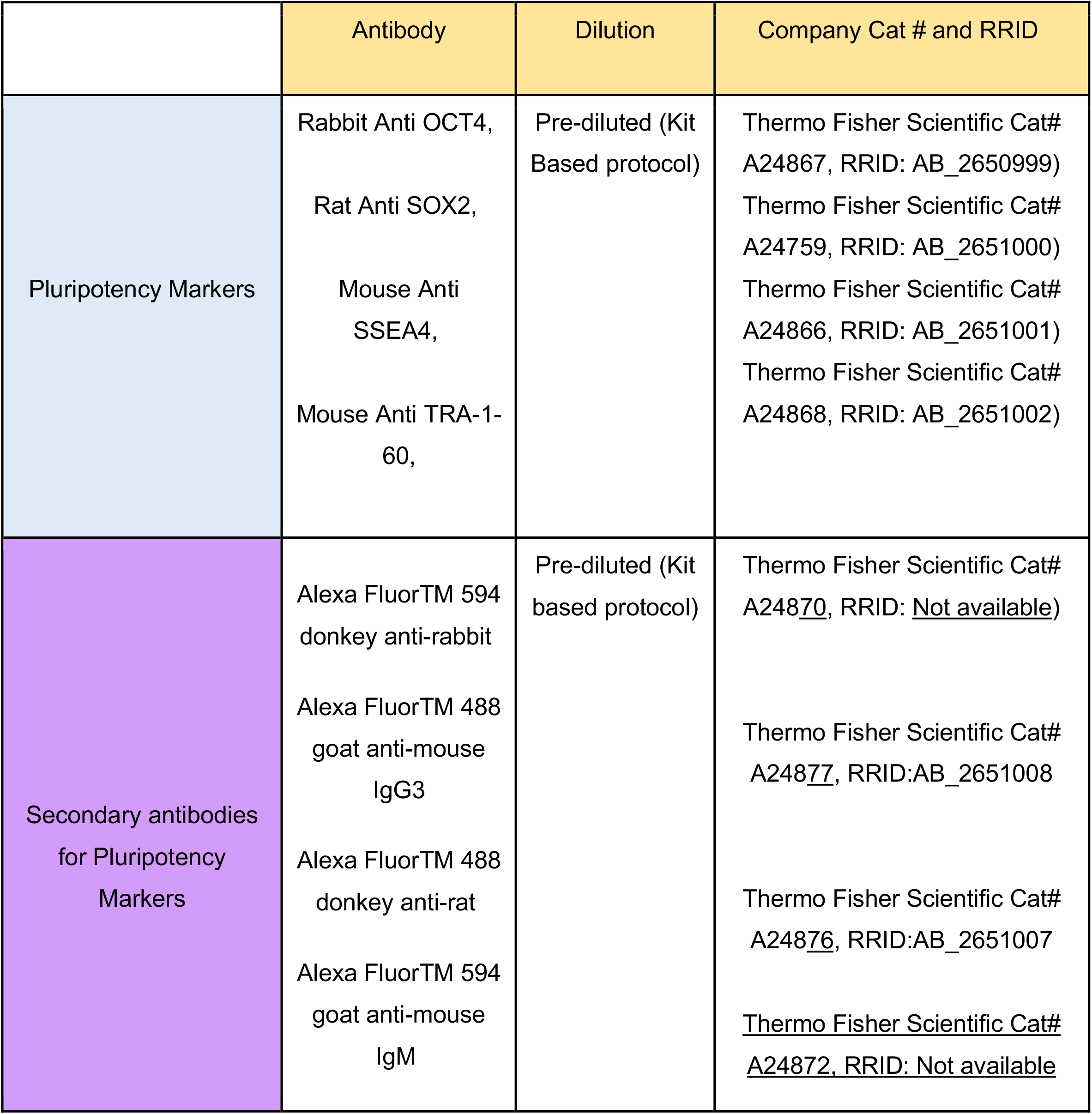
Antibodies for immunofluorescence staining of hIPSCs for pluripotency markers.

**Table 3.**
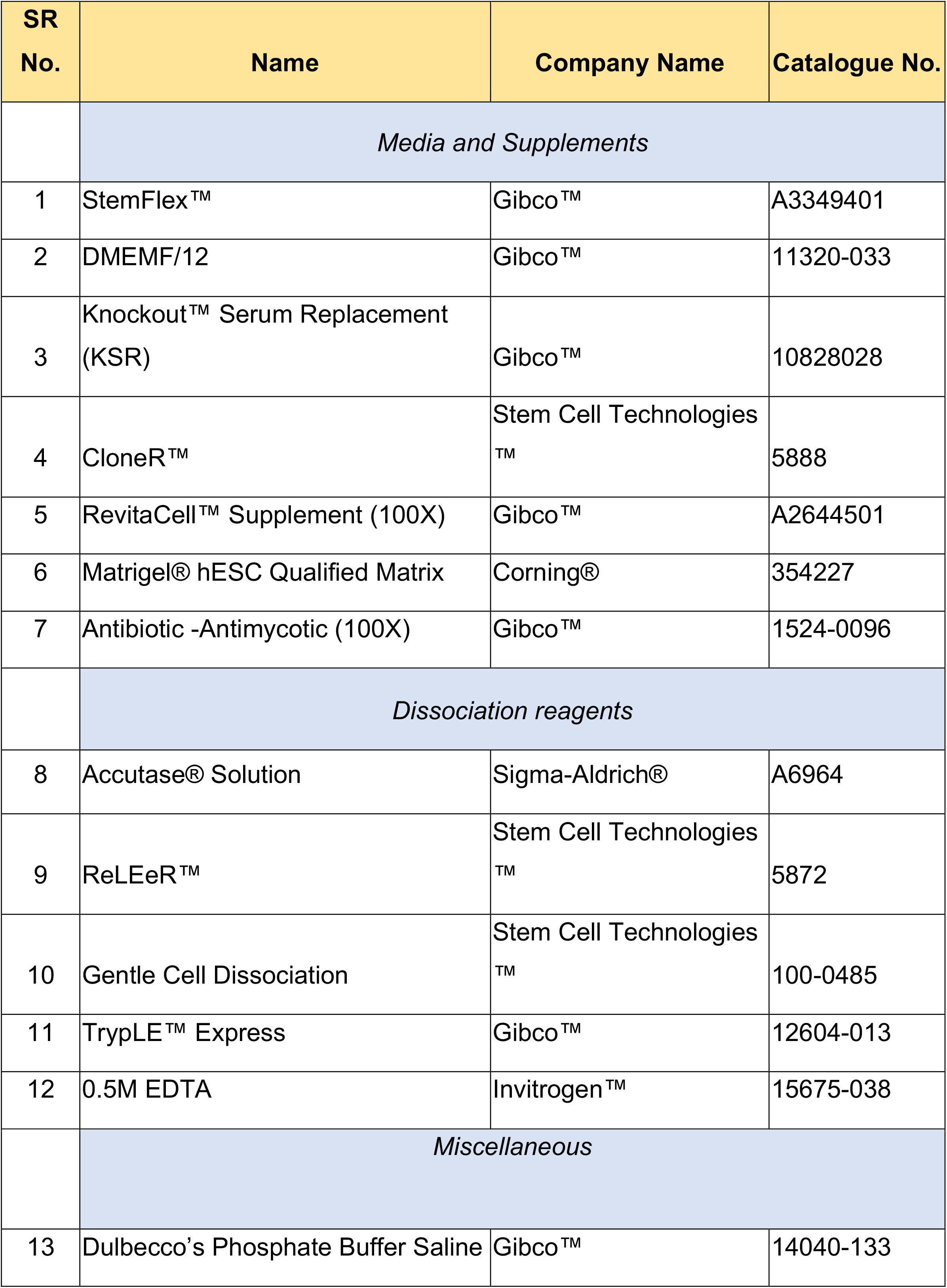
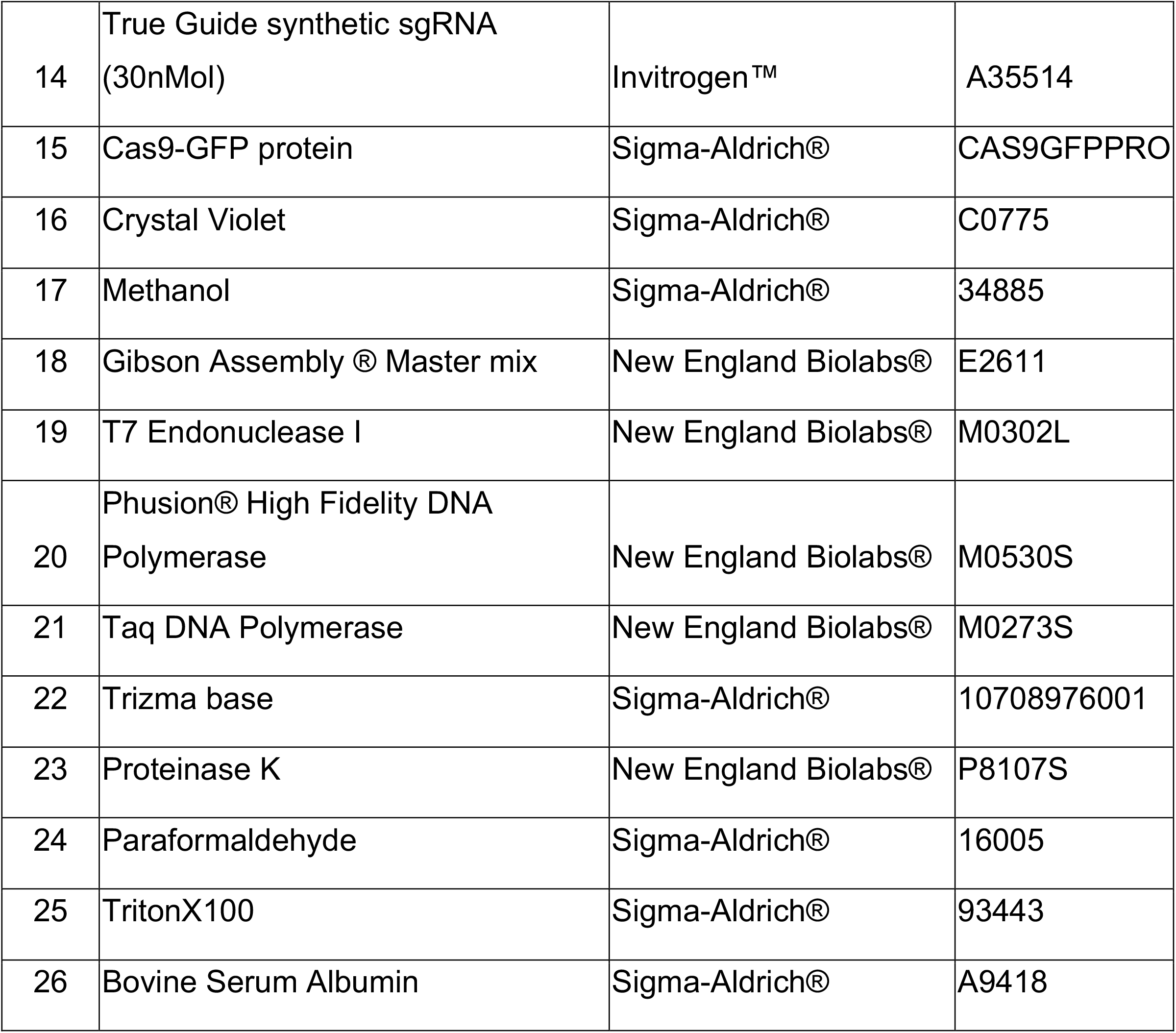
Material List.

**Table 4.**
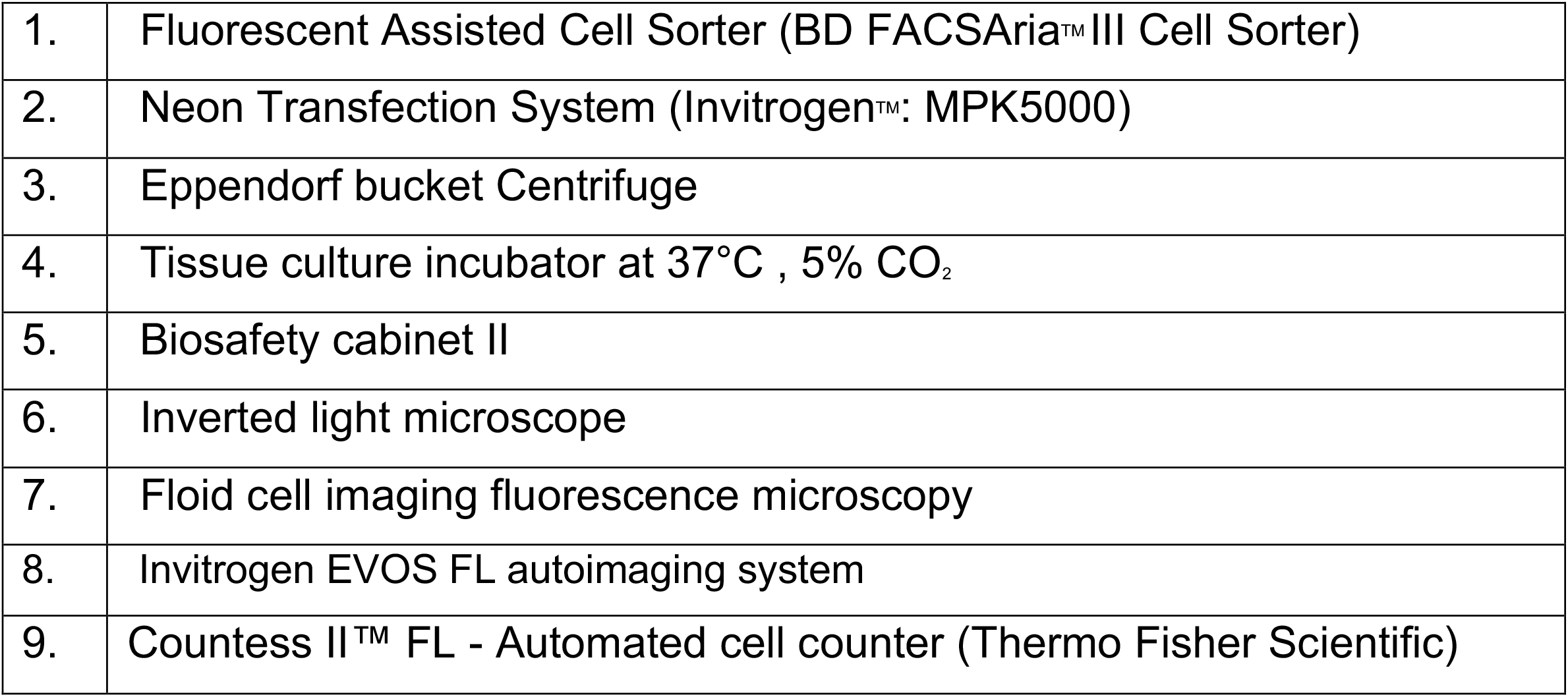
Equipment List.

**Supplementary Figure 1.**
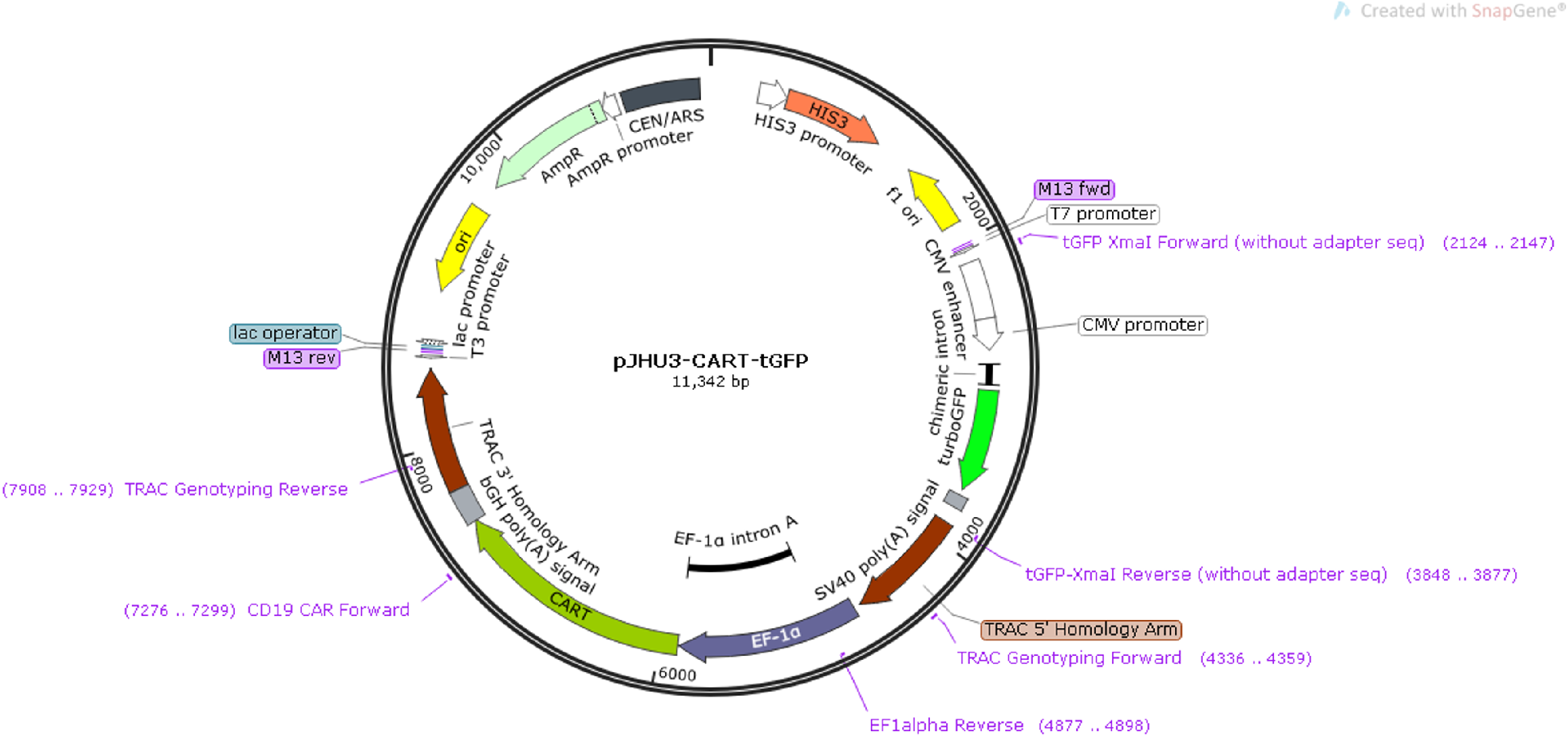
pJHU3-CART-tGFP donor vector map for KI experiment.

**Supplementary Figure 2.**
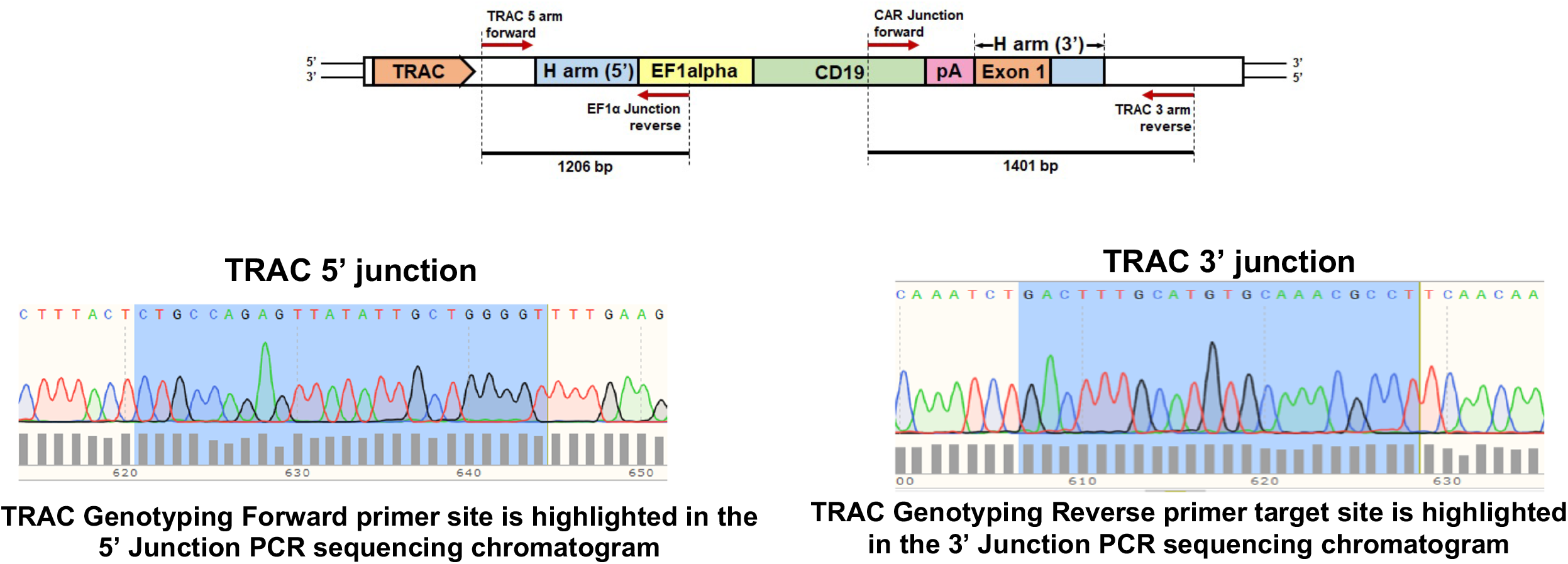
Sequence traces of 5’ and 3’ Knock-in sites for confirmation of targeted insertion in hiPSC.

